# MOLECULAR AND MORPHOMETRIC APPROACHES IN RECENTLY RADIATED SPECIES: *Inga subnuda* SALZM. EX BENTH AND *Inga vera* WILLD. (LEGUMINOSAE, MIMOSOID CLADE)

**DOI:** 10.1101/2025.02.01.636048

**Authors:** Clara da Cruz Vidart Badia, Flávia Cristina Pinto Garcia, Larissa Areal Muller, Jèferson Nunes Fregonezi

## Abstract

The species-rich genus *Inga* (Leguminosae) presents ca. 300 species widespread throughout the Neotropics and is recognized by its recent and rapid diversification. Forty-eight species of *Inga* are endemic of the Brazilian Atlantic Forest. Among them, *Inga subnuda*, with two subspecies: subsp. *subnuda*, which occurs from the state of Paraíba to Rio de Janeiro; and subsp. *luschnathiana*, which occurs from Espírito Santo to the state of Santa Catarina. Both subspecies occur in sympatry in the southeastern region and share floral and leaf characters, which hampers the morphological delimitation. The co-occurrence of *Inga vera* subsp. *affinis*. with both *I. subnuda* subspecies results in intermediate morphologies between *I. vera* subsp. *affinis* and *I. subnuda* subsp. *luschnathiana*, making the distinction between the latter species even harder. We sampled 94 individuals from 8 natural populations and evaluated morphological characters previously described as distinctive among the subspecies of *I. subnuda* in addition to others not measured yet. We used 84 plastid (*trnD-trnT* spacer) and 58 nuclear (ITS 1 and 2) sequences to characterize the phylogenetic relationships between the taxa. The results obtained point out *I. subnuda* subsp. *subnuda* as a more structured taxon in relation to the other subspecies, whilst *I. subnuda* subsp. *luschnathiana* and *I. vera* subsp. *affinis* constituted a cohesive group. The apportionment of haplotypes differed between the markers used, thus evincing a retention of ancestral polymorphism between the subspecies, given their recent diversification. This paper explores different lines of evidence, thus contributing to the delimitation of these species. Demographic and biogeographic scenarios are also discussed. A new status of for the taxon currently circumscribed as *Inga subnuda* subsp. *luschnathiana* is suggested.

---

The legume genus *Inga* Mill (Leguminosae) is represented by ca. 300 woody species and is widespread throughout the Neotropics (Richardson et al. 2001). *Inga* presents 132 species spread in Brazil, of which 51 are endemic of the Atlantic Forest (Pennington 1997). The high species richness of *Inga*, added to its short generation time contributes to the species richness in the Neotropical rainforests (Richardson et al. 2001).

The speciation of the genus is estimated to have occurred in the past 4.3 million years, with a major diversification in the last 2 million years, when the Neotropical climates were less stable (Richardson et al. 2001). The monophyly of the genus is supported by molecular and morphological synapomorphies (Richardson et al. 2001, Brown 2008), but its infrageneric phylogenetic relationships remains poorly understood. Among the endemic species within the brazilian Atlantic Forest there is *Inga subnuda* Salzm. ex Benth., which is currently circumscribed as presenting two subspecies: *Inga subnuda* Salzm. ex Benth. subsp. *subnuda*. and *Inga subnuda* subsp. *luschnathiana* (Benth.) T.D.Penn. The first is distributed geographically from Paraíba State (northeastern Brazil) to Rio de Janeiro State (southeastern Brazil), occurring in sympatry with subsp. *luschnathiana* in Espírito Santo, Minas Gerais and Rio de Janeiro States. The latter subspecies occurs also in São Paulo, Paraná and Santa Catarina States (Garcia and Fernandes 2015).

Sympatrically to *I. subnuda* subspecies there is *Inga vera* subsp. *affinis* (DC.) T.D.Penn., which occurs from Colombia to Uruguay. The distinct combinations of continuous varying characters defines the delimitation within *Inga* species (Bentham 1845, Pennington 1997), but in particular cases the delimitation is hampered by characters that can be shared between distinct *taxa* (Bentham 1845, Pennington 1997).

*Inga vera* subsp. *affinis* is particularly hard to tell apart from other *I. vera* subspecies, and also from *I. subnuda* subsp. *luschnathiana* (Bentham 1845, Pennington 1997, Garcia 1998). Visually, *I. vera* subsp. *affinis* presents more morphological similarities with *I. subnuda* subsp. *luschnathiana* than between both *I. subnuda* subspecies.

In order to evaluate the morphological similarities and bioclimatic suitability areas distributed among *I. subnuda* subspecies, Castro (2018) performed morphometric analysis and ecological niche modeling for these *taxa*. Floral and vegetative characters discriminate both subspecies, which is expected since both subspecies are easily distinguished visually. Ecological niche overlap was found in the southeastern region for *I. subnuda* subspecies. Both subspecies presented expansion of its suitability areas during the last interglacial (LIG). Regarding the last glacial maximum, a retraction of suitability areas was noticed for *I. subnuda* subsp. *subnuda*, thus corroborating the refugia described for Bahia and Pernambuco regions (Carnaval and Moritz 2008). The interspecific relationships concerning *I. subnuda* and *I. vera* subsp. *affinis*, however, remains unknown.

The genetic studies performed so far concerning *Inga* do not include *I. subnuda* or *I. vera* subsp. *affinis*, although other subspecies of *I. vera* are occasionally sampled (Richardson et al. 2001, Kursar et al. 2009, Neto et al. 2014).

Because *Inga* constitutes a recently radiated genus (Richardson et al. 2001), some of the plastid markers traditionally used are not informative at the species level (e.g. intergenic spacers *trnL-trnF, trnH-psbA, trnS-trnG* and íntron *trnL*). Also, neither next generation sequencing methods provide fully resolved phylogenies (Nicholls et al. 2015). In this study, we used molecular and morphometric approaches in order to investigate whether *I. subnuda* subspecies are close enough to be considered as subspecies. We characterized the diversity and looked for genetic structure between natural populations of both subspecies sampled along a representative range. Also, morphometric analysis was performed using characters previously noted as useful to discriminate both subspecies. We included *I. vera* subsp. *affinis* as an in-group in both analysis in order to obtain a higher accuracy of the possible evolutionary relationships between these *taxa*. Further discussions concerning morphological similarities, phylogenetic relationships with sister groups and biogeographic history within *I. subnuda* are provided in this work.

## Materials and Methods

### Study Area

This study was carried out through field collections of populations in the Atlantic Forest, from Bahia to Santa Catarina states, between 14°19’S 39°1’W and 26°4’S 48°36’W, along altitudes from 0 to 826 m above sea level (Fig. 1). We accessed Restinga, Alluvial and Submontane Dense Ombrophilous Forests of Atlantic Domain in order to collect *I. subnuda* subsp. *subnuda* and *I. subnuda* subsp. l*uschnathiana*, since this species occurs more often in these vegetal formations. *Inga subnuda* subsp. *subnuda* populations located at Bahia and northern Espírito Santo states are included in Af and Aw climates respectively, according to Köppen’s climate classification (tropical without dry season and tropical with dry winter (Alvares et al., 2013). The climatic classification of the localities of natural populations distributed along Espírito Santo, Minas Gerais, Rio de Janeiro, Paraná and Santa Catarina states are included in the humid subtropical zone (Cfb, Cwb, Cfb and Cfa, respectively). Specimens sampled in Restinga were only found in fragments, evidencing that the anthropic occupation of the coast may be a threat to populations of both subspecies.

**FIG. 1.**
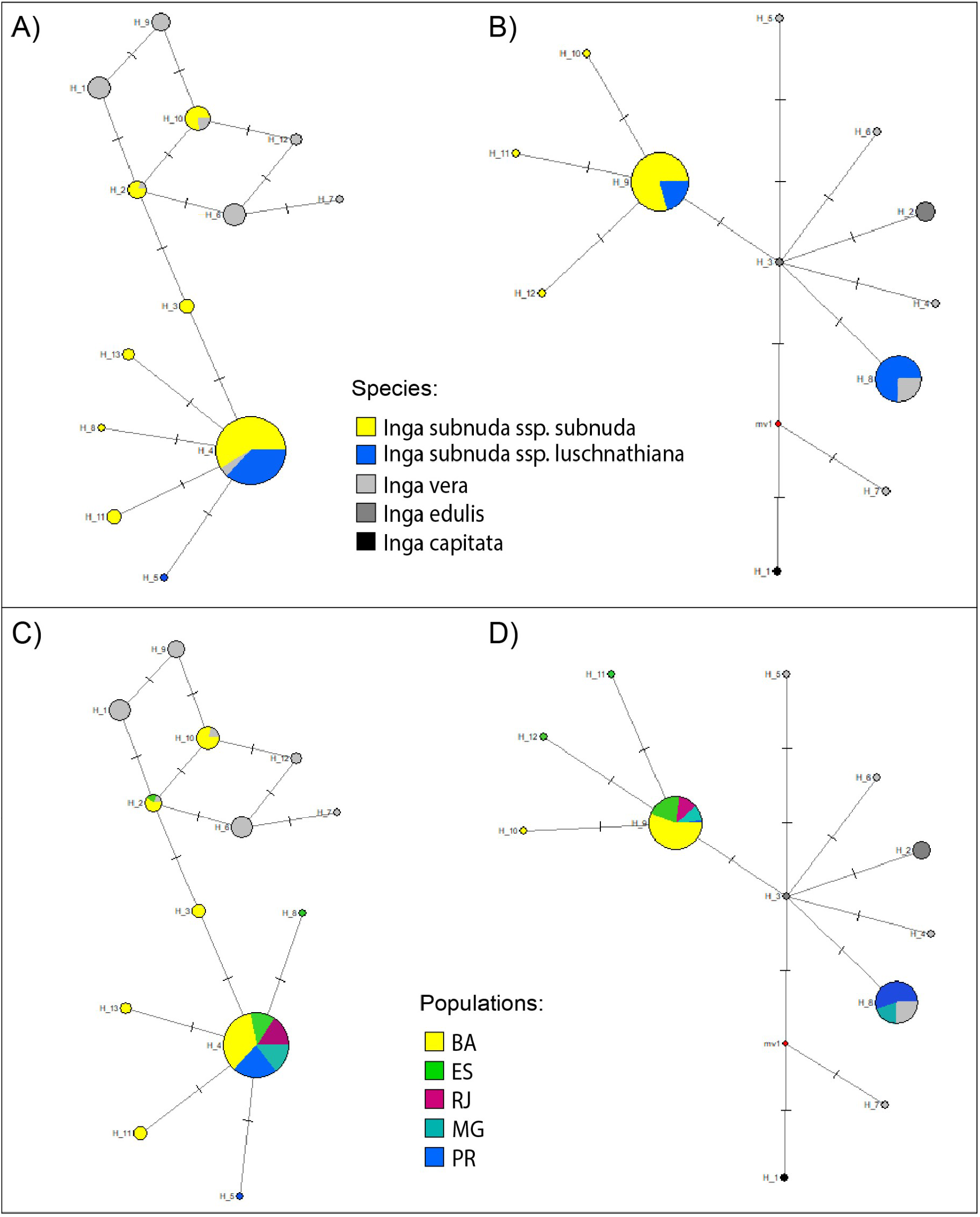
Geographical distribution of *Inga subnuda* subsp. *subnuda* and *Inga subnuda* subsp. *luschnathiana* in Brazil based on herbarium records (coloured ellipses). Sampling localities (coloured dots) and corresponding population codes are also indicated.

### Specimen Sampling

Literature and herbarium specimens occurrence records were consulted to conduct field excursions aiming to comprise the widest part of the subspecies distribution, as well as its morphological variation. A total of 94 individuals from 8 distinct locations were collected, covering a significant part of the species’ distribution. The specifications of localities, codes and respective number of individuals per population are summarized in Table 1 and detailed in Appendix 1. Fresh leaves of each specimen were collected and stored in silica-gel, and all sampled specimens were herborized, identified by comparison with herbarium material and consultation with specialized literature and with specialists (Pennington 1997, Garcia 1998). Vouchers were deposited in the VIC Herbarium, Botany Department, Federal University of Vicosa, Brazil.

**TABLE 1.** Population codes (POP.CODES), number of individuals sampled for genetic analysis (T*rnD*-*trnT*, ITS and concatenated markers *TrnD-trnT* + ITS), number of individuals used in morphometric analyses (MORPHO.ANALYSIS) and locations of *Inga subnuda* subsp. *subnuda* (ISS), subsp. *luschnathiana* (ISL) and *Inga vera* subsp. *affinis* (IVA) natural populations sampled. Geographical coordinates of populations are provided in Appendix 1.

| SPECIES/SUBSPECIES | POP.<br>CODES | MORPHO.*TrnD*-<br>ANALYSIS | *TrnD*-<br>*trnT* | ITS | *TrnD*-<br>*trnT* + ITS | LOCATIONS |
| --- | --- | --- | --- | --- | --- | --- |
| *Inga subnuda* subsp. *subnuda* | BA-North | 23 | 20 | 9 | 9 | Areia (PB), Ilhéus, Serra Grande and Itacaré |
|  | BA-South | 15 | 5 | 12 | 4 | Ilhéus, Canavieiras and Una |
|  | ES-North | 0 | 4 | 4 | 4 | Nova Viçosa, Itaúnas and São Mateus |
|  | ES | 2 | 3 | 0 | 0 | Domingos Martins and Santa Teresa |
|  | RJ | 5 | 5 | 5 | 5 | Búzios |
| *Inga subnuda* subsp. *luschnathiana* | ES | 3 | 4 | 1 | 1 | Santa Teresa |
|  | MG-Brig | 12 | 4 | 4 | 4 | Araponga, Ervália and Viçosa |
|  | MG-JF | 6 | 5 | 1 | 0 | Juiz de Fora, Lima Duarte and Santa Bárbara do Monte Verde |
|  | PR | 18 | 16 | 7 | 7 | Guaratuba, Itapoá (SC), Matinhos and Morretes |
| *Inga vera* subsp. *affinis* | BA-South | 1 | 0 | 0 | 0 | Una |
|  | MG-Brig | 3 | 1 | 1 | 1 | Araponga and São Gonçalo do Rio Preto |
|  | RJ | 2 | 2 | 2 | 2 | Petrópolis |
|  | PR | 4 | 3 | 1 | 1 | Guaratuba, Itapoá (SC) and Morretes |
| OVERALL |  | 94 | 72 | 47 | 38 |  |

The sampled populations comprised typical morphologies of the subspecies towards north for *I. subnuda* subsp. *subnuda* and south for *I. subnuda* subsp. *luschnathiana*, as well as areas in which both subspecies occurs simpatrically. Therefore, intermediate morphologies that eventually might occur have also been sampled. Individuals of *Inga vera* subsp. *affinis* (DC.) T.D.Penn. were sampled along the distribution range covered for both *Inga subnuda* subspecies and used as in-group in morphometrics and molecular analyses.

### DNA Extration, Amplification and Sequencing

Populations of *Inga subnuda* subspecies and *Inga vera* subsp. *affinis*, used here as an in-group, were sampled along its distribution range for genetic studies (number of individuals and respective populations are described in table 1). Genomic DNA was extrated from silica gel-dried leaves using the CTAB method described by Roy et al. (1992) and electrophoresis was performed in 1% agarose gel in order to check DNA integrity. NanoDrop® Spectrophotometer (Thermo Scientific, Wilmington, DE, USA) were also performed to quantify concentrations and access the purity of DNA samples. The chloroplast intergenic spacer *trnD*-*trnT* and the nuclear ribosomal internal transcribed spacers (ITS1 and 2) were chosen with the purpose of investigating genetic diversity and population structure of *Inga subnuda* subspecies. These markers presented higher number of polymorphic sites compared to other genomic regions used in previous studies for *Inga* species (Richardson et al. 2001, Coley et al. 2005, Dexter et al. 2010, Simon et al. 2011). In preliminary tests we also evaluated the utility of the intergenic spacers *trnH*-*psbA, trnS*-*trnG* and *trnL*-*trnF*, as well as íntron *trnL* (Taberlet et al. 1991, Shaw et al. 2005), commonly used in similar studies, which did not show genetic variation in the sample set composed for *Inga subnuda* subspecies and *Inga vera* subsp. a*ffinis*.

The following primers were used for the amplification of *trnD*-*trnT* region: *trnD*2 (Simon et al. 2011; GTGTACAGCATGCATATTCTTACG) and *trnT*ggu (Simon et al. 2011; CTACCACTGAGTTAAAAGGG). For ITS region amplification, we used the primers ITS4 (White et al. 1990; TCCTCCGCTTATTGATATGC) and ITS5A (Stanford et al., 2000; CCTTATCATTTAGAGGAAGGAG). The internal primers ITS2 (White et al., 1990; GCTGCGTTCTTCATCGATGC) and ITS3 (White et al., 1990; GCATCGATGAAGAACGCAGC) were used in addition to ITS4 and ITS5A in order to obtain more reliable sequences. PCR reactions for *trnD*-*T* region were conducted in a total volume of 50 µL, in a final concentration of 1× buffer, 300 µM each dNTP (Promega), 0,2 µM each primer, ½U of Go*Taq* polymerase (Promega) and 10-20 ng of template DNA. The conditions for ITS marker were conducted in a total volume of 50 µL containing: 1× buffer, 200 µM each dNTP, 0,2 µM each primer, 2% Dimethyl sulfoxide (DMSO), 1U of Go*Taq* polymerase and 0.1-10 ng template DNA. Amplification conditions for *trnD*-*T* were: 94 ºC for 5 min, 30× 94 ºC for 45 s, 55 ºC for 1 min, 72 ºC for 1 min and a final extension of 5 min at 72 ºC; and for ITS: 94 ºC for 3 min, 30× 94 ºC for 1 min, 55.5 ºC for 1 min, 72 ºC for 1 min and a final extension of 10 min at 72 ºC. PCR produts were purified using ExoSAP protocol (provided by ThermoFisher) and sequenced by Macrogen (Seoul, South Korea).

### Sequence Analyses and Genetic Diversity

The sequences were analyzed for both forward and reverse primers using Chromas software v.2.6.6 (available at *https://technelysium.com.au/wp/chromas/*). In addition to *Inga subnuda* subspecies and *Inga vera* subsp. *affinis* sampled for this work, we retrieved *Inga capitata* Desv., *Inga edulis* Mart. and *Inga vera* Willd. sequences from GenBank database in order to add sister groups in the dataset. The DNA alignment was generated using Muscle algorithm (Edgar 2004) in MegaX software (available at *https://www.megasoftware.net/*; Kumar et al. 2018) and edited manually. Vouchers and respective GenBank accession numbers are provided in Appendix 1.

Molecular analyses were performed using 84 sequences for *trnD*-*trnT* plastid marker and 58 sequences for ITS marker. Haplotypes of cpDNA were estimated using DnaSP v. 6.12.03 software (Rozas et al. 2017). In order to ensure that heterozigous sites were included correctly in haplotypes reconstruction, nuclear sequences were analysed using PHASE software version 2.1.1 (Stephens et al. 2001). Genetic diversity parameters such as haplotype diversity (*H*) and nucleotide diversity (π) were retrieved from Arlequin 3.5.2.2 (Excoffier and Lischer 2010). Polymorphic regions observed in poly-A/T regions usually found among cpDNA sequences were not included in the analysis, since its homologies cannot be equatelly accessed (Aldrich et al. 1988). Because adjacent nucleotides tends to evolve together, insertion/deletion mutational events containing more than one base pair (bp) were considered as a single mutation (Simmons and Ochoterena 2000, Hartl and Clark 2010).

### Phylogenetic Relationships

Evolutionary relationships among haplotypes were accessed through median-joining network method, as implemented in Network v.10 software (available at https://www.fluxus-engineering.com/sharenet.htm).

Phylogenetic reconstructions were performed using Bayesian inference in BEAST v.1.8.0 (Suchard et al. 2018). ITS and cpDNA datasets were treated as independent partitions, and priors were assigned to each partition separately. A Yule Tree Prior model was used in BEAST, since it is most suitable for trees describing the relationships between individuals from different species, considering speciation processes through birth of new lineages. We adopted the HKY substitution model, as it is the most appropriate evolution model inferred by Bayesian Information Criteria in JmodelTest software v.2.1.7 (Darriba et al. 2012). For molecular dating we assumed the lognormal relaxed molecular clock and nucleotide substitution rates previously used for plastid and nuclear markers for *Inga* species (ITS: 2.34 x 10^-9^ s/s/y; *trnD*-*trnT*: 1.30 x 10^-9^ s/s/y; Richardson et al. 2001). As the standard deviation was not available, we assigned the value of 0.2 x 10^-9^ so that the normal distribution would stay within the expected for the generation time of *Inga*. The Markov Chain Monte Carlo (MCMC) was run for 20 million generation, being 20% of the trees discarded as burn-in. In order to ensure that the effective sample size was higher than 200, we checked the results in Tracer software v.1.7.1 in BEAST package. We accessed the node ages and posterior probabilities in TreeAnnotator v.1.8, also included in BEAST package. The trees obtained were designed in FigTree v.1.3.1 (Rambaut 2010).

A second reconstruction was performed in BEAST using a coalescent-based species-tree analysis, with both molecular markers with unlinked gene-trees. Other parameters and priors were adjusted as described above. This approach sets species as priors in order to check for clear distinction between the *taxa*.

A phylogenetic rescontruction based on Maximum Likelihood method was also performed in MegaX software, with substitution models and other parameters determined by jModeltest software as described previously. Branch supports were obtained by bootstrap method (Felsenstein 1985) with 100 replicates.

### Genetic Structure and Demographic Analysis

The effective populations’ size over time was accessed using Bayesian Skyline Plot analysis (Drummond et al. 2005), available in BEAST, using the same priors previously described.

In order to evaluate genetic structure between the populations sampled, we concatenated the individuals sampled for both markers, totaling 34 individuals, being 22 of subsp. *subnuda* and 12 of subsp. *luschnathiana*, distributed among 7 populations (Table 1, detailed in Appendix 1). Since it was not our aim to sample populations of *I. vera* subsp. *affinis*, only isolated individuals as an in-group, this species was not included in this analysis. We analyzed this dataset using the Bayesian Clustering Algorithm embedded in software BAPS (Bayesian Analysis of Population Structure)(Corander and Marttinen 2006, Corander et al. 2008, Tang et al. 2009), adopting the admixture clustering model and assuming 5 replications for each population. Were tested hypotheses for the existence of genetic clusters from K= 2 to K=7, without assuming any previous subspecies or populations as priors.

### Morphometric Analysis

Ninety four individuals with fully developed leafs and flowers or fruits were selected and measured prior to statistical analyses. In this dataset, 35 individuals presented only flowers and 55 individuals presented only legumes. The remaining specimens (4 individuals) presented both structures when collected. In order to include a higher number of specimens and to standardize as much as possible the divergent availability of fertile traits per specimen, we considered the length of the pedicel of the fruit for the specimens which presented both flowers and fruits. The individuals of *Inga subnuda* subsp. *subnuda* and *Inga subnuda* subsp. *luschnathiana* were distributed between 8 populations and the individuals of *Inga vera* subsp. *affinis* were considered as supplementary data (for more information, see Table 1). Because only the data of 2 individuals of *Inga subnuda* subsp. *subnuda* and 3 of subsp. *luschnathiana* was available for the same location, the population “ES”, from Espirito Santo, comprised both *Inga subnuda* subspecies. Pennington (1997) and Garcia (1998) were used for initial determinations, as well as consultation with specialist. We chose 4 morphological traits that obtained statistically significant results by Castro (2018) (WAW, WE, LE and CP) and additional 3 vegetative characters (WP, NP and NR) according to personal observation of what could contribute to separate both subspecies (Table 2). The quantitative morphological structures were measured using digital caliper (precision 0.01 mm) and respective descriptions are presented in detail in Table 2. Eventual missing data was substituted for the mean value of the character of the belonging population of a given specimen. Principal Component Analysis was performed in RStudio (Rstudio Team, 2015) using a matrix of qualitative and quantitative characters (considering leaf and flower/fruit variables) of both *I. subnuda* subspecies. In order to reduce discrepancy in measurements, quantitative data was standardized using logarithmic transformation. The dataset was checked for the assumptions of normality (Shapiro et al. 1968) and homocedasticity (Levene 1960), and the difference between the unrelated groups were accessed for PC1 using ANOVA (Kaufmann and Schering 2014)

**TABLE 2.** Morphological characters used for morphometric analyses of the two subspecies of *Inga subnuda* Salzm. ex Benth and *Inga vera* subsp. *affinis* and respective descriptions.

| CHARACTER | DESCRIPTION |
| --- | --- |
| WP | Winged petiole (presence/ absence) |
| NP | Lenght of naked petiole (mm) |
| NR | Lenght of naked portion of proximal rachis (mm) |
| WAW | Width of distal apex wing (mm) |
| WE | Width of stipules (mm) |
| LE | Lenght of stipules (mm) |
| LP | Lenght of pedicel (mm) |

## Results

### Genetic Diversity

Considering *trnD*-*trnT* plastid marker, 37 sequences were obtained for subsp. *subnuda*, 29 for subsp. *luschnathiana* and 18 of sister groups (*I. vera* (11), *I. capitata* (1) and *I. edulis* (6)), totaling 84 sequences with 1405 base pairs long (bp) in the alignment. Twelve haplotypes were identified based on 13 polymorphic sites, being 4 parcimoniously informative. Haplotype (*H*) and nucleotide (*π*) diversities for this marker were, respectively, 0.6182 ± 0.1643 and 0.0009 ± 0.0007 for *I. vera;* 0.4433 ± 0.0685 and 0.0006 ± 0.0005 for subsp. *luschnathiana;* and 0.1577 ± 0.0802 and 0.0001 ± 0.0002 for subsp. *subnuda* (Table 3).

**TABLE 3.** Haplotype (*H*) and nucleotide (*π*) diversities and respective standard deviation (±) of each species considering plastid and nuclear markers. The diversities for concatenated sequences from both markers is provided as “overall”. IV = *Inga vera*; ISS = *Inga subnuda* subsp. *subnuda*; ISL = *Inga subnuda* subsp. *luschnathiana*.

| SPECIES | HAPLOTYPE DIVERSITY ($H$) | NUCLEOTIDE DIVERSITY ($\pi$) |
| --- | --- | --- |
|  | PLASTIDIAL |  |
| IV | 0.6182 ± 0.1643 | 0.0009 ± 0.0007 |
| ISS | 0.1577 ± 0.0802 | 0.0001 ± 0.0002 |
| ISL | 0.4433 ± 0.0685 | 0.0006 ± 0.0005 |
| SPECIES | NUCLEAR |  |
| IV | 0.8805 ± 0.0360 | 0.0030 ± 0.0012 |
| ISS | 0.4977 ± 0.0753 | 0.0016 ± 0.0012 |
| ISL | 0.0769 ± 0.0697 | 0.0001 ± 0.0003 |
| SPECIES | OVERALL |  |
| IV | 0.9103 ± 0.0485 | 0.0008 ± 0.0006 |
| ISS | 0.6203 ± 0.0800 | 0.0006 ± 0.0005 |
| ISL | 0.4502 ± 0.1120 | 0.0004 ± 0.0003 |

For the nuclear marker (ITS), a total of 58 sequences were obtained, being 30 sequences of subsp. *subnuda*, 13 of subsp. *luschnathiana* and 15 of *I. vera, containing* 648 base pairs in the DNA alignment. The 9 polymorphic sites observed (5 parcimoniously informative) were distributed among 13 haplotypes. Four heterozygous sites were identified within PHASE results. The number of nucleotide substitutions obtained for both *I. subnuda* subspecies were relatively low, which is consistent for *taxa* that have recently radiations (Richardson et al. 2001). Haplotype (*H*) and nucleotide (*π*) diversities for this marker were, respectively, 0.8805 ± 0.0360 and 0.0030 ± 0.0012 for *I. vera;* 0.0769 ± 0.0697 and 0.0001 ± 0.0003 for subsp. *luschnathiana;* and 0.4977 ± 0.0753 and 0.0016 ± 0.0012 for subsp. *subnuda* (Table 3).

Considering sequences sampled for both markers, 41 sequences were analyzed, being 23 of subsp. *subnuda*, 11 of subsp. *luschnathiana* and 7 of *I. vera*, totaling 2053 bp. *Inga vera* presented haplotype (*H*) and nucleotide (*π*) diversities of 0.9103 ± 0.0485 and 0.0008 ± 0.0006 respectively; subsp. *luschnathiana* obtained 0.4502 ± 0.1120 and 0.0004 ± 0.0003; and subsp. *subnuda*, 0.6203 ± 0.0800 and 0.0006 ± 0.0005 (Table 3). The difference found between chloroplast and nuclear marker is expected, since the first only inherit mutations from maternal lineage and the latter can accumulate mutations from both parental lineages, in addition to suffer recombination events.

### Phylogenetic Relationships

Both *trnD-T* and ITS haplotype networks exhibit few mutations that separate the subspecies of *Inga subnuda*, as well as few mutations between the other species, considered more taxonomically distant. In addition there are haplotype sharing between species and subspecies in the analyzed samples (Fig. 2A, B).

**FIG. 2.** Haplotype median-joining network obtained for ITS and *TrnD-trnT* markers. Each circle represents a haplotype, and circle sizes are proportional to the overall frequency of the haplotypes. The colors within each circle indicates taxonomic entities or populations. The cross marks represent mutational differences between haplotypes. The ougroups *Inga vera, Inga edulis* and *Inga capitata* are represented in grey tones and are not included in the population analysis since sequences were retrieved from GenBank (Appendix 1). A) Network obtained for ITS marker representing *Inga subnuda* subspecies and the sister groups. B) Network obtained for *trnD-trnT* marker representing *Inga subnuda* subspecies and the sister groups. C) ITS network representing the distribution of haplotypes within and between the sampled populations. D) *TrnD-trnT* network representing the distribution of haplotypes within and betweem the sampled populations.

As *Inga* is noted as a recently radiated group (Richardson et al. 2001), it is possible to identify within the *trnD*-*T* haplotype network (Fig. 2B) that the central haplotype H3 from sister group *I. edulis is* separated from *I. subnuda* and *I. vera* subspecies for one mutational event and the outer group *I. capitata* (H1) for two mutations. These results highlights that even morphologically distinct and well delimited species presented little genetic differences between them in plastid haplotype network.

Haplotype sharing is observed between *I. subnuda* subsp. *luschnathiana* and *I. vera* (H8 in Figure 2B) as well as between both *I. subnuda* subspecies (H9 in Fig. 2B).

The ITS haplotype network, in which samples of *I. edulis* and *I. capitata* were not included, there is also little genetic distance. Haplotype sharing between *I. subnuda* subsp. *subnuda* and *I. vera* (H2 and H10 in Figure 2A) and between the two subspecies of *I. subnuda* and *I. vera* can also be noted (H4 in figure 2A).

In both haplotype networks, generated from plastid and nuclear regions (Figs. 2A, B), several haplotypes are present in few individuals, and they are connected to one more frequent haplotype, separated by a single mutation each. In ITS network (Fig. 2A) H4 haplotype is more frequent and is connected by one mutational event to H3, H5, H8, H11 and H13 haplotypes. Likewise, the H9 haplotype of *trnD-T* network (Fig. 2B) is more frequent, connected by one mutation to H10, H11 and H12 haplotypes, present in few sampled individuals. This “star-like” pattern shown in both networks have peripheral haplotypes mainly related to individuals belonging to *I. subnuda* subsp. *subnuda*. The exception is the H5 haplotype in the ITS network (Fig. 2A), belonging to one individual from *I. subnuda* subsp. *luschnathiana*.

Distinct haplotype relationships patterns among *I. subnuda* subspecies and *I. vera* can be observed when comparing the networks from both markers. In ITS network (Fig. 2A), *I. vera* haplotypes are shared (H2, H10) and more related to haplotypes belonging to *I. subnuda* subsp. *subnuda*. On the other hand, the *trnD-T* network shows one *I. vera* haplotype (H8) shared with *I. subnuda* subsp. *luschnathiana* individuals, which reflects the similar morphology of these *taxa*.

When haplotypes are marked by sampled locations for two subspecies in both haplotype networks, there is no clear pattern between haplogroups and populations (Figs. 2C, D). However, it can be noted that those peripheral, low frequent haplotypes present in both networks belong mainly to individuals of *I. subnuda* subsp. *subnuda* from northernmost geographic distribution (BA and ES populations, yellow and green colors in Figs. 2C, D). The individuals located in central regions of the sampled distribution (RJ and MG populations, purple and light blue colors in Figs. 2C, D) have mainly shared haplotypes that are in high frequency (H4 in Fig. 2C; H9 in Fig. 2D). The exceptions are H5 haplotype of ITS (Fig. 2C), and H8 haplotype of *trnD-T* (Fig. 2D), which belong to individuals from the south of the sampled distribution (PR population, dark blue color in Figs. 2C, D).

Phylogenetic reconstructions recovered trees with very low node support values. The Maximum Likelihood analysis showed low support (<60) for all nodes, while the Bayesian method indicated only two groups with reasonable support values. Both reconstructions revealed the same topologies and relationships between individuals. The tree obtained by Bayesian Method is shown in Fig. 3. The calculated ages for supported groups, as well as the age for the most recent common ancestor for the entire sample are also shown in Fig. 3. As expected, the ages for these clusters were very low, with an average age of about 511.000 years for the common ancestor of *I. capitata, I. edulis, I. subnuda* and *I. vera*.

**FIG 3.**
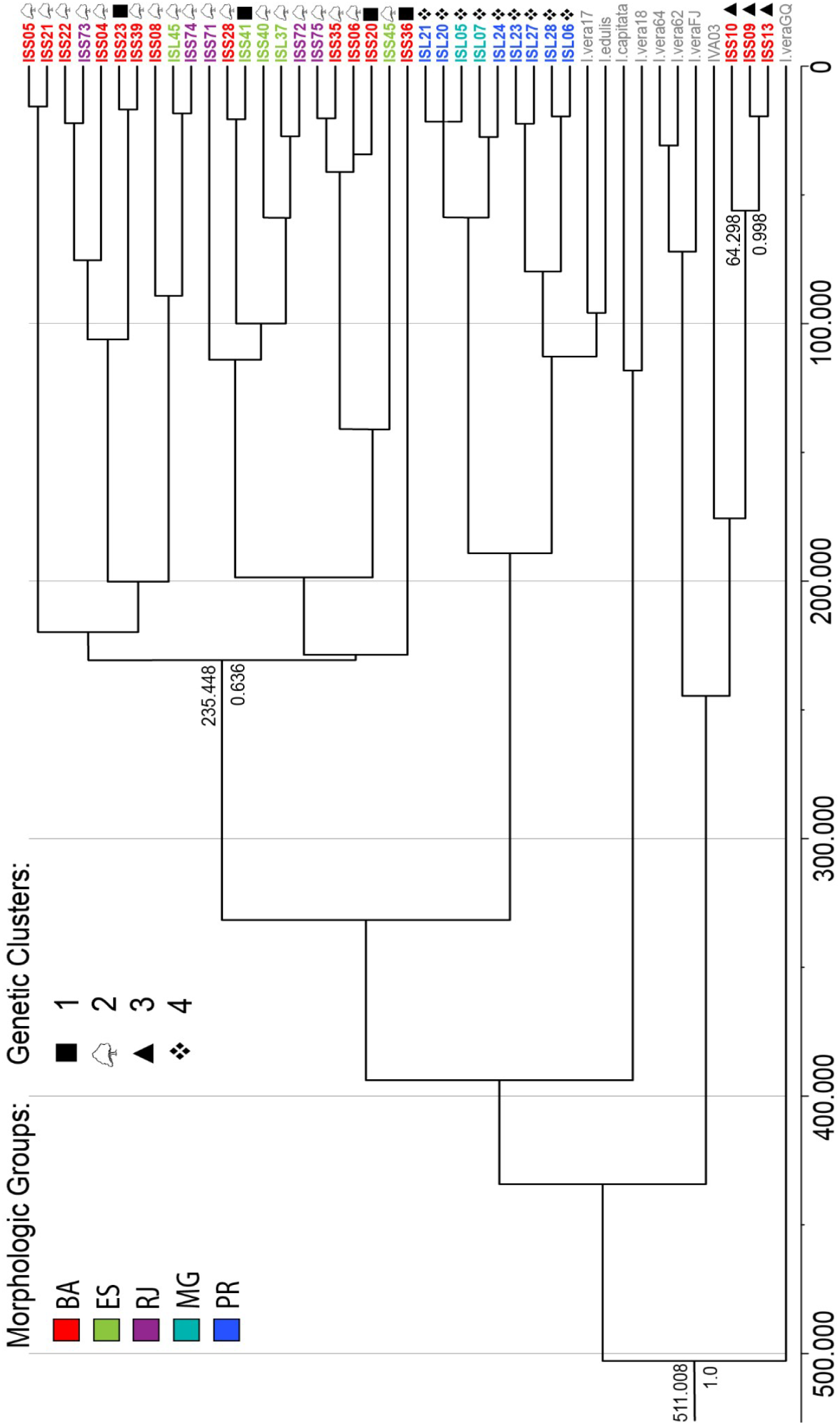
Bayesian phylogenetic tree indicating evolutionary relationships between *Inga subnuda* subspecies considering the morphological groups by morphometric analysis (colored by morphological groups). The posterior probabilities (PP>0.6) are represented below branches and ages (in years) are indicated above branches. A temporal scale is represented below the phylogenetic topology. The genetic structure provided by BAPS are indicated by distinct symbols beside each individual. The sister groups *Inga vera, Inga edulis* and *Inga capitata* are indicated in grey.

Regarding the topology obtained through Bayesian inference, the two supported groups can be identified (Fig. 3). The first one (posterior probability, PP = 0.636, age of 235.448 years) comprises most part of *Inga subnuda* subsp. *subnuda* specimens sampled in BA, ES and RJ populations. The second group (PP = 0.998, age of 64.298 years) includes three individuals from *I. subnuda* subsp. *subnuda*, being closest to *I. vera* samples than to subsp. *luschnathiana*. Another interesting grouping (wich are not supported by PP values) includes mainly individuals of *I. subnuda* subsp. *Luschnathiana* from MG and PR populations.

### Genetic Structure and Demographic Analysis

The results provided by BAPS (Genetic groups marked by colors in Fig. 4 and indicated by symbols in the phylogenetic tree of Fig. 3) pointed 4 genetic clusters, of which one gathers both *Inga subnuda* subspecies from all the localities, mainly those in which the subspecies occur in sympatry (cluster 2 marked in green in Fig. 4). Cluster 1 assembled *Inga subnuda* subsp. *subnuda* from BA and ES populations (yellow colors in Fig. 4) whilst cluster 3 presented only *I. subnuda* subsp. *subnuda* from Northern Bahia localities (red color in Fig. 4).

**FIG. 4.**
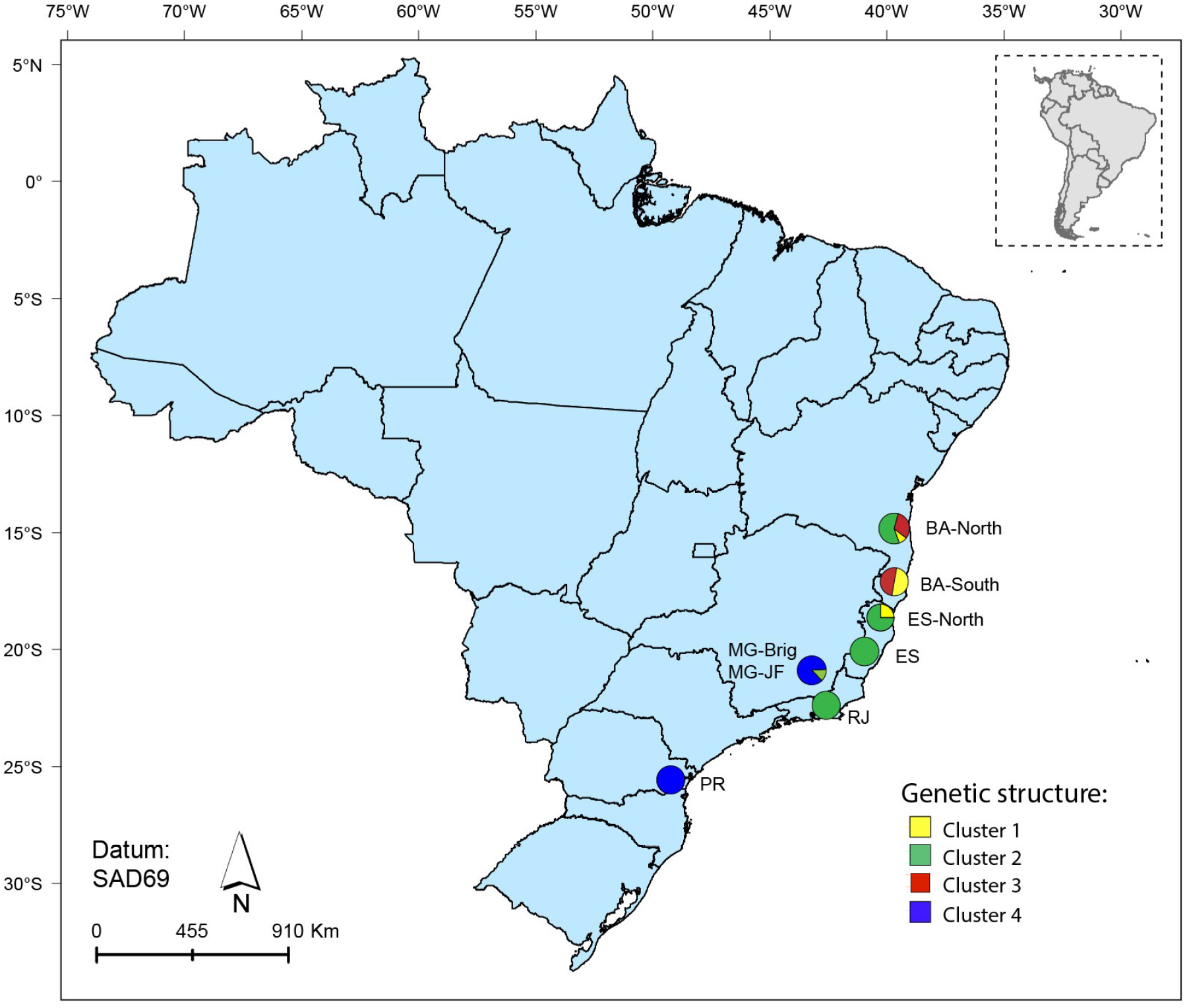
Result provided by BAPS (Bayesian Analysis of Population Structure) using Bayesian Clustering Algorithm. Each color represents one of the four genetic clusters defined by the analysis. Pie charts colored according to the proportion of each genetic clustering within and between populations. Populations names are indicated beside each pie chart.

Even though *I. subnuda* subsp. *luschnathiana* seem to be not as structured as *I. subnuda* subsp. *subnuda*, the cluster 4 (marked in blue in Fig. 4) grouped individuals of this subspecies located in central regions of the distribution (MG populations) and southward locations (PR population).

Three genetic clusters appear to be geographically structured (1, 3 and 4 in Fig. 4), and the individuals of mixed localities comprised in cluster 2 (18 individuals) seem to lack of a clear pattern that reflects its geographical distribution. *I. subnuda* subsp. *subnuda* seems to be more spatially structured if compared with *I. subnuda* subsp. *luschnathiana*.

The Bayesian Skyline Plot Analysis indicates that *I. subnuda* subsp. *subnuda* may have slightly increased in effective population size in the last 2 thousand years ago (Kya), compared to *I. subnuda* spp. *luschnathiana*, which did not show changes in the average population size over time for this analysis. However, this results do not present statistical support due to its high confidence intervals displayed in population sizes plots.

### Morphometric Analysis

The PCA results shows strong relationships between the variables (Fig.5). 79.30% of the morfological variability can be explained by two principal components. The first principal component (PC1) explains 63.90% of the data variability, and PC2 explains 15.40%. The variables that contributed the most to PC1 were the lenght of naked portion of proximal rachis (NR; 88.07% of contribution), width of distal apex wing (WAW; 65.24%) and length of pedicel (LP; 58.50%). The variable that contributed the most to PC2 was lenght of naked petiole (NP), with 60.32% of contribution. Out of the three distinctive morphological characters, “lenght of naked portion of proximal rachis”, “width of distal apex wing” are conspicuous and very useful to distinguish *I. subnuda* subsp. *subnuda* visually. Due to the highest level of information carried by PC1, hereafter we will only refer to this dimension for further discussion, since this axis seem to be the one that possesses a substantial information. The results are shown in figure 5.

**FIG. 5.**
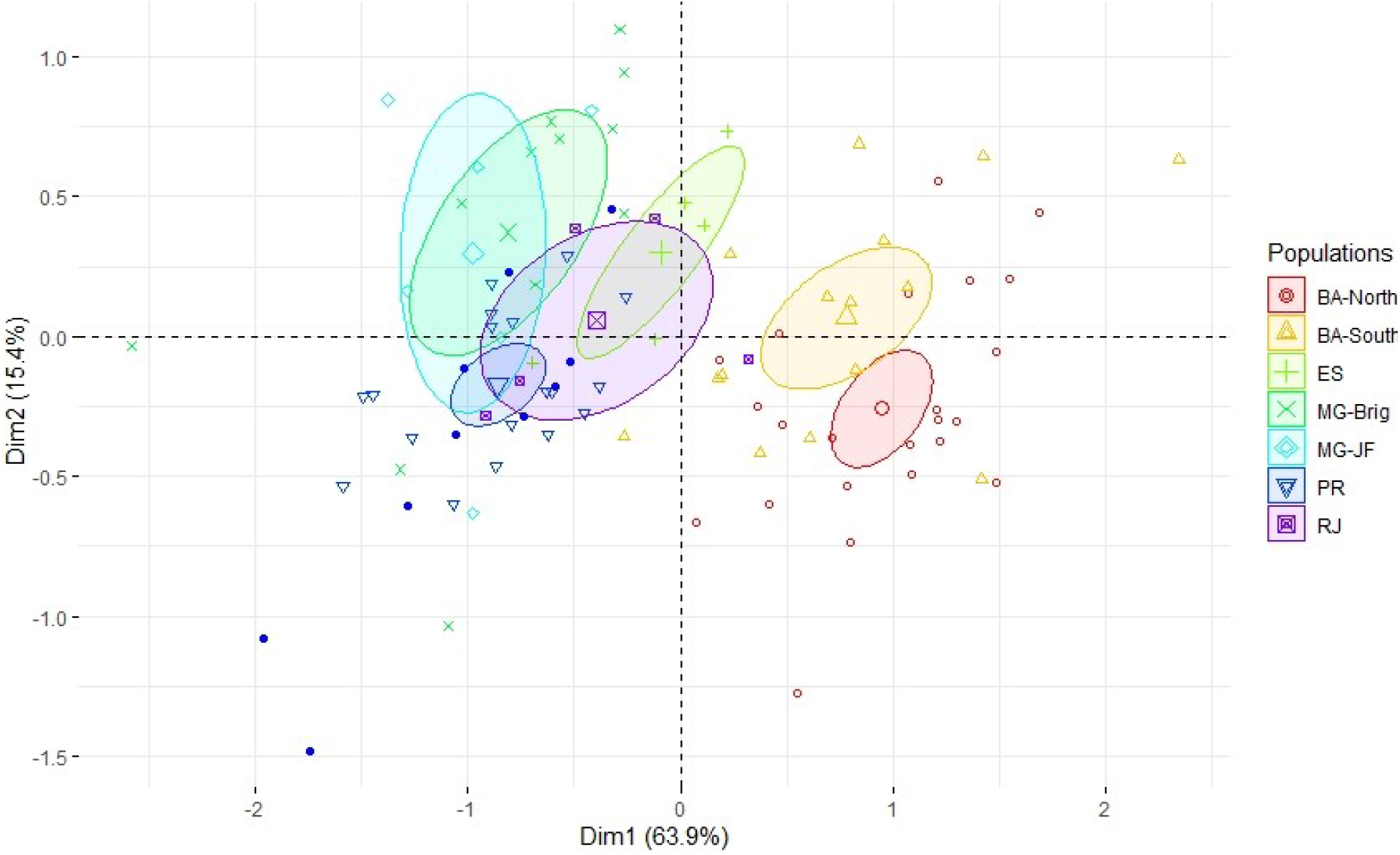
Graphic derived from Principal Component Analysis considering principal components 1 (“Dim1” in the PCA plot) and 2 (“Dim2”). The individuals of *Inga subnuda* subspecies included in the analysis are described by their populations codes (Table 1, Appendix 1). *Inga vera* subsp. *affinis* was included as supplementary data and is represented by blue dots. The bigger symbols centralized in the ellipses indicates the mean value that represents each population.

ANOVA results applied in PC1 shows that two main groups are present in the dataset, corresponding with the PCA groups. One group contains populations of Bahia, which comprises only *I. subnuda* subsp. *subnuda* individuals, and the other group contains all populations of south and southeastern Atlantic Forest regions, which includes *I. subnuda* subsp. *luschnathiana* and only one individual of *I. subnuda* subsp. *subnuda* from Espirito Santo State. Moreover, supplementary individuals of *Inga vera* subsp. *affinis* were grouped with *I. subnuda* subsp. *luschnathiana’*s individuals.

## Discussion

### Genetic Nature and Species Proximity in Inga

Estimating genetic properties within *taxa* that radiated recently can be challenging. *Inga* stands out as an example of rapid and recent diversification taxon comprised among the species-rich rainforests of the Neotropics (Richardson et al. 2001). That said, inferences concerning the genetic nature in a species level, such as phylogenetic relationships and populational structure, can be gruelling and are commonly poorly supported (Richardson et al. 2001, Nicholls et al. 2015). The genus is estimated to have arisen in the last 4.3 Mya, and more intensively diversificating in the Pleistocene (ca. 2 Mya)(Richardson et al. 2001). The phylogenies obtained so far show low resolution between the species, presenting short branches with low support value, thus presenting poor genetic differentiation between the *taxa*.

The main drivers of this rapid radiation seem to be ecological and climatic. The rapid evolution of leaf chemical responses against herbivores is suggested as playing an important role in competition, therefore contributing to *Inga* diversity and abundance among the Neotropical rainforests (Kursar et al. 2009). The climatic oscillations during the Pleistocene affected mostly the Atlantic Forest species’ distribution (Carnaval and Moritz 2008). The historically stable areas among the glacial and interglacial periods seem to have contributed to a higher genetic pool over time (Richardson et al. 2001, Carnaval and Moritz 2008).

A phenomena commonly found among recent and rapid diversificated *taxa* is described as retention of ancestral polymorphism in phylogenetic and populational analysis. During incomplete lineage sorting, species gene pools share genetic information from ancestral species after speciation, by retention of ancestral polymorphism succeeded by fixation of different alleles in the descendant lineages (Maddison 1997, Charlesworth et al. 2005). It can result in haplotype sharing among different recent radiation species, as well as conflicting genealogies when comparing different genes or genome regions.

This seems to be the most plausible scenario for the subspecies studied in this work. The phylogeny obtained did not present high support values (Fig. 3). In both reconstructions (Bayesian, Fig. 3 and Maximum Likelihood analysis, not shown), there was no high confidence values for the groups found, even though the topologies and the relationships between the sampled individuals are similar. Besides, the species-tree phylogenetic reconstruction (data not shown) resulted in low ESS values, indicating that there are no totally isolated genetic lineages among the sampled individuals when compared the separate trees for each data set (nuclear and plastid).

The haplotype networks (Fig. 2) present high number of shared haplotypes between different *Inga* species, notably among subspecies of *I. subnuda* and *I. vera* subsp. *affinis*. Moreover, nuclear haplotype networks relationships (Figs. 2A, C) differed from those obtained for the plastidial region (Figs. 2B, D), a pattern often described among other plant species (Rieseberg and Soltis 1991, Soltis and Kuzoff 1995, Tsutsui et al. 2009) which can indicate two possible scenarios: recent radiation or hybridization between well-established species. It is very difficult to distinguish incomplete lineage sorting and hybridization because both can give rise to patterns of shared genetic diversity.

ITS haplotype network points out that *I. subnuda* subsp. *subnuda* is closely related to *I. vera* subsp. *affinis* instead of *I. subnuda* subsp. *luschnathiana*, as indicated in the plastid networks. This incongruence could be explained by hybridization events, in which the nuclear DNA was inherited from both *I. vera* subsp. *affinis* and *I. subnuda* subsp. *subnuda*, and the cpDNA uniparentally inherited from *I. vera* subsp. *affinis*. Over the next generations of hybrids, the influence of *I. subnuda* subsp. *subnuda* in the nuclear haplotypes among the hybrids reduced, whereas the frequency of *I. vera* subsp. *affinis’* cpDNA was the same since the introgression first occurred. The newly arisen lineage, currently corresponding to *I. subnuda* subsp. *luschnathiana*, would then share plastidial haplotypes with *I. vera* subsp. *affinis* and present distinct and/or intermediate morphologies, closest to *I. vera* subsp. *affinis*.

Since the *taxa* in question are part of a recently radiated genus and the time of divergence was not enough for the *taxa* to speciate and generate hybrids, a complete introgression process is less likely to have occurred. The results obtained here evidences that the three *taxa* are recent: low values of nucleotide diversity; lack of population structure separating both *I. subnuda* subspecies and *I. vera* subsp. *affinis*; lack of hypothetical haplotypes (median vectors) among the plastid networks – which would be an evidence of divergent *taxa*; and low posterior probability support and short branches among phylogeny trees.

### Morphological and Genetic Relationships

Morphometric analysis results shows that *Inga subnuda* subsp. *subnuda* differs from subsp. *luschnathiana* and *I. vera* in the northern-most populations sampled, which comprises mainly Bahia locations (for more information, see table 1 and Fig. 5). The Principal Component Analysis (PCA) evidences that *I. subnuda* subsp. *subnuda* populations of Bahia presents a clear and highly supported distinction, indicated by PC1 in Fig. 5, from *I. subnuda subsp. luschnathiana* and *Inga vera* subsp. *affinis*.

The morphological group obtained for subsp. *subnuda* is congruent with the genetic patterns obtained by BAPS genetic clustering and haplotype networks. Individuals of this morphological group are placed in genetic clusters 1 and 3 defined by BAPS (Fig. 4 and indications in Fig. 3). Individuals from cluster 1 share the same haplotype regarding the *trnD-T* network (Haplotype 9; Fig. 2B, D) and the same two haplotypes within ITS networks (Haplotypes 2 and 4; Fig. 2A, C). A similar pattern is represented in the networks for cluster 3, where two out of the three individuals share the *trnD-T* haplotype 9. Considering the ITS network, haplotype 10 comprise all the individuals of this cluster, being one of them also represented within haplotype 11. These individuals also constitute a cohesive group considering the Bayesian inference, in which the posterior probability value is the highest observed for all samples (0.998, Fig. 3, indicated by triangles in the terminals).

The genetic cluster 4 defined by BAPS has individuals of *I. subnuda* subsp. *luschnathiana* from MG and PR populations, which shared the same *trnD-trnT* and ITS haplotypes (H8 and H4 respectively, Fig. 2). Additionally, one sample is represented by an additional ITS haplotype (H5), differing from the others towards south, where this individual is located. This clustering is not supported by the phylogenetic tree (Fig. 3), but is highly supported by Principal Component Analysis (79.30%, Fig. 5), where a clear distinction is found between southern and southeastern *I. subnuda* subsp. *luschnathiana* and northern *I. subnuda* subsp. *subnuda* populations.

In the regions in which the three *taxa* co-occur, the morphological distinction is not so clear. PCA results points out that southeastern populations (ES and RJ) of *I. subnuda* subsp. *subnuda* are morphologically mixed with *I. subnuda* subsp. *luschnathiana* populations (Fig. 5) and the morphological distribution of *I. vera* subsp. *affinis* is associated with this mixed group. BAPS analysis placed these individuals in genetic cluster 2, presenting the majority of individuals from BA, RJ and ES populations, with some *I. subnuda* subsp. *luschnathiana* individuals from ES population (Figs. 3 and 4). Among these results, the individuals of *I. subnuda* subsp. *subnuda* sampled RJ population were assembled with the northern-most locations above-mentioned for this subpsecies.

Despite this admixture of some individuals in group 2 indicated by BAPS and the overlapping of morphologies in the central region of the geographical distribution, the discrimination for northern *I. subnuda* subsp. *subnuda* populations is quite clear by morphological (Fig. 5) and genetic results (Figs. 3 and 4).

The subspecies of *I. subnuda* are generally well delimited morfologically from other species of the genus, but *I. subnuda* subsp. *luschnathiana* and *I. vera* subsp. *affinis* are particularly hard to tell apart (Bentham 1845, Pennington 1997, Garcia 1998). Our results shows that south (PR) and southeastern (MG) populations are genetically related to *I. vera* subsp. *affinis*, as shown in *trnD-trnT* network (Fig. 2B, D). These *I. subnuda* subp. *luschnathiana* populations are grouped according to BAPS analysis (cluster 4 in Fig.4, with indication in Fig. 3) presenting the same distribution as in the PCA plot. PCA results shows clearly a higher morphological similarity between *I. subnuda* subsp. *luschnathiana* and *I. vera* subsp. *affinis* than between *I. subnuda* subspecies.

### Demographic and Biogeographic Patterns

Bayesian skyline plot results suggests that *I. subnuda* subsp. *subnuda* has slightly increased its effective population size in the last 2 thousands of years (kya). Moreover, the higher number of peripheral haplotypes observed for *I. subnuda* subsp. *subnuda* (represented by a “star-like” pattern for both genomic regions in haplotype Networks, Fig. 2) and the higher overall haplotype diversity in comparison to *I. subnuda* subsp. *luschnathiana* are also evidences that point out a recent population expansion. This information is consistent with the results obtained by Castro (2018), in which ecological niche modeling regarding bioclimatic data shows an increase in suitability areas for *Inga subnuda* subsp. *subnuda* in this time period.

In contrast, the same pattern cannot be fully applied to *I. subnuda* subsp. *luschnathiana*, once there is no star-like pattern or further evidence that there was an increase in effective population size. In fact, *I. subnuda* subsp. *luschnathiana* haplotypes were more cohesive, presenting only one derived haplotype towards south. The derived haplotype 5 of ITS network (Figs. 2A, C) might be an evidence that this taxon is also under expansion, but in a smaller and more recent scale. This isolated haplotype could represent some sort of populational growth, but may also be a reflect of a subsampled population.

Both *Inga subnuda* subspecies were found as spatially structured towards north and south, and in the southeastern region where both subspecies occur in sympatry the pattern becomes unclear. The lack of structure within the central Atlantic Forest, as well as the shared haplotypes between southeastern populations (Fig. 2C, D) and the lack of mutational events within these locations support the hypothesis that both *Inga subnuda* subspecies may have arisen within the southeastern region, therefore expanding their ranges towards north and south.

Among the Brazilian Atlantic Forest, historically stable areas known as refugia were described by Carnaval and Moritz (2008). Some of the described refugia are located in the central corridor comprised in Bahia and Espirito Santo and also in the State of Pernambuco. The presence of historical forest stability in *I. subnuda* subsp. *subnuda* northern-most occurrence could have contributed to a retention in its populational range and haplotype diversity in the glacial periods (reflecting cohesive and well-defined groups, such as clusters 1 and 3 defined by BAPS, in Fig. 4, as well as the well-supported clade found in the phylogenetic tree in Fig. 3).

In the southern portion of the Brazilian Atlantic forest, unstable climatic stretches of rainforest are noted instead of small refugia (Leite et al. 2015), a data supported by the glacial Quaternary environmental conditions predominantly characterized by cooler climate, significantly lower sea levels and reduced precipitation. The suitability areas obtained for *I. subnuda* subsp. *luschnathiana* by Castro (2018) corroborate this hypothesis, as well as the lower diversity of this subspecies for the southern locations. The haplotype sharing between *I. subnuda* subsp. *luschnathiana* and other taxa from the southeastern region, in addition to its derived haplotype noted for the southern Atlantic Forest region, support the scenario that *I. subnuda* subsp. *luschnathiana* arose in southeastern region and then differentiated towards south.

### Taxonomic Considerations and Conclusions

The *taxa* currently circumscribed as subspecies of *Inga subnuda* were initially defined by Bentham (1845, 1876) as *I. subnuda* and *I. luschnathiana* Benth. Based on observation of specimens from Rio de Janeiro, Espírito Santo and Bahia States, Pennington (1997) identified intermixed leaf and floral traits, therefore considering *I. luschnathiana* as conspecific of *I. subnuda*. With this circumscription, *I. subnud*a was regarded occupying two extremes of a range of variation: pediceled flowers, terete petiole and rachis, legume margins keeled when immature, faces almost completely covered by the expanded margins and mature legume more or less cylindrical corresponding to *I. subnuda* subsp. *subnuda;* and short pediceled or sessile flowers, winged rachis, legume margins not keeled, faces exposed and mature legume more or less quadrangular corresponding to *I. subnuda* subsp. *luschnathiana* (Pennington 1997, Garcia 1998).

Although this taxonomic treatment apparently helped to disentangle the continuous variation of characters commonly observed between *Inga* species (Pennington 1997), the difficulty of distinguishing these two subspecies have been discussed (Garcia 1998, Castro 2018). Aiming to better understand the delimitation between these *taxa*, Castro (2018) applied morphometric analysis and ecological niche modeling to a dataset comprised by herbarium specimens. The morphological leaf traits found as distinctive between both *Inga subnuda* subspecies were the length of petiole, length of stipule and width of apex wing, whilst the distinctive floral characters were the length of floral rachis, length of stamens, length of peduncle and length of pedicel.

The ecological niche modeling results obtained by Castro (2018) corroborated the distinction between both subspecies. The southern coastal plain and northeast area of the Atlantic Forest was identified as suitable for *I. subnuda* subsp. *subnuda*, whereas for *I. subnuda* subsp. *luschnathiana*, suitability areas were found throughout the southeast coastal region between Rio de Janeiro and northeast Santa Catarina States. Moderate ecological niche overlap was found in the southeastern region of Atlantic Forest. Furthermore, expansion in both *I. subnuda* subspecies suitability areas was described for the last interglacial (LIG). During the last glacial maximum (LGM), expansion was recorded for the suitability areas of *I. subnuda* subsp. *luschnathiana*, whereas *I. subnuda* subsp. *subnuda* presented retraction in its suitability areas for the same period.

Despite the distinction between *Inga subnuda* subspecies became clearer after Castro (2018) results, no studies regarding the difficulties discussed by authorities of the genus (Bentham 1845, Pennington 1997, Garcia 1998) in distinguishing morphologically *Inga subnuda* subsp. *luschnathiana* from *Inga vera* subsp. *affinis* was performed until now.

The morphological distinction within *I. subnuda* subspecies is quite clear. *Inga subnuda* subsp. *luschnathiana* is easily distinguished from *I. subnuda* subsp. *subnuda* by its faces exposed and margins striate, length of naked portion of proximal rachis and length of pedicel. The conundrum is that *I. subnuda* subsp. *luschnathiana* is frequently confused with *I. vera* subsp. *affinis* when one come across an intermediate morphology, even though they are described under distinct taxonomic entities. When analyzing typical morphologies, the calix open in bud, narrower and smaller tube calix, and shorter lobes calyx presented in *I. subnuda* subsp. *luschnathiana* make this distinction possible. In spite of that, intermediate morphologies of the latter two subspecies are almost impossible to place (Bentham 1845, Pennington 1997, Garcia 1998).

In this work, we assembled a new dataset comprised of specimen sampled along the distribution range including natural populations of both *I. subnuda* subspecies and some individuals of along the distribution range *I. vera* subsp. *affinis*, so that the morphological and genetic relationships between these *taxa* could be enlightened. We accessed the same morphological traits previously described as exploitative to distinguish *I. subnuda* subspecies (Castro 2018) and added some other characters (Table 2) to check which ones would better explain the similarities and differences between those groups.

We found that the morphological similarities between *I. vera* subsp. *affinis* and *I. subnuda* subsp *luschnathiana* observed in field collections and described among other works (Bentham 1845, Pennington 1997, Garcia 1998) are statistically supported (Fig. 5). Therefore, a closer phylogenetic relationship between these taxonomic entities is evinced (Figs. 2, 3, 4 and 5; see Results section for details). These results indicate that the current taxonomic treatment proposed by Pennington (1997) is not accurate, since it does not reflect the relationship between the *taxa*.

Both morphological and genetic evidences (Figs. 2, 3, 4 and 5; see previous sections for more details) point out that *I. subnuda* subsp. *subnuda* constitutes a more cohesive and well delimited group in comparison to *I. subnuda* subsp. *luschnathiana* and *I. vera* subsp. *affinis*, even though the results show a possible scenario of retention of ancestral polymorphism commonly observed among recently radiated genus (Steiner et al. 2012, Goetze et al. 2017).

To confirm this pattern, sampling of the northern-most part of *I. subnuda* subsp. *subnuda* distribution, spread above Bahia state, would be required. Furthermore, throughout the field excursions and observations of herborized material, we perceived that the trichome density is different between *I. vera* subsp. *affinis* and *I. subnuda* subsp. *luschnathiana*. Trichomes are a very informative character among Leguminosae (Çildir et al. 2012, Grohar et al. 2016) and other taxa (Ndukwu and Agbagwa 2006, Tschan and Denk 2012, Seyedi and Salmaki 2015). Although studies regarding this character are not performed very often for *Inga* species, after careful study of several specimens we believe that it could contribute to distinguish more accurately these two *taxa*.

Chemocoding proved to be useful to understand the taxonomic relationships within the genus *Inga*, specially when neither morphology or DNA sequencing are able to answer how the species are related to each other (Endara et al. 2018). Accordingly, further studies using this method would contribute to the understanding concerning the species of this work.

Hence, according to results found in this study, we reinforce the circumscription proposed by Castro (2018), in which he suggest to retrieve the circumscription made by Bentham (1845, 1876). In this way, each subspecies of *I. subnuda* would return to the status of species.

## Acknowledgments

The authors thanks the Botany Graduate Program (PPGBot) and the Federal University of Viçosa (UFV) for providing the necessary structure to carry out this work. We thank the National Council for Scientific and Technological Development (CNPq - Ministry of Science and Technology - Brazil) for research scholarship, the Coordination for the Improvement of Higher Education Personnel (CAPES-PROAP, Brazil) and Minas Gerais State Agency for Research and Development (FAPEMIG) for research fundings (CRA-APQ-017003-17). Special thanks to Sara Abduani Brum and Cecília Vieira Miranda, for their help in the field excursions. We also appreciate the assistance given by Dr. Marcelo Gomes Marçal Vieira Vaz (Plant Growth Unit – UFV), whom provided resources and supervision in the laboratory of Molecular Biology. This work was led by the first author as Master’s student, as part of the requirements for obtaining the academic degree on *Magister Scientiae* at UFV, under the supervision of Dr. Jéferson Nunes Fregonezi. Funding was partially provided by UFV.

## Author Contributions

BADIA, C.C.V. Collected field and laboratory data, analyzed data, wrote the manuscript and provided the manuscript figures.

GARCIA, F.C.P. Idealized the research, guided the field collections and confirmed specimen identification.

Muller, L.A. Contributed with morphometric data analysis.

FREGONEZI, J.N. Idealized, led and guided the research. Contributed to data collection through field excursions and guided the collection of laboratory data, as well as data analysis and the writing of the manuscript.

---

**Appendix 1.**
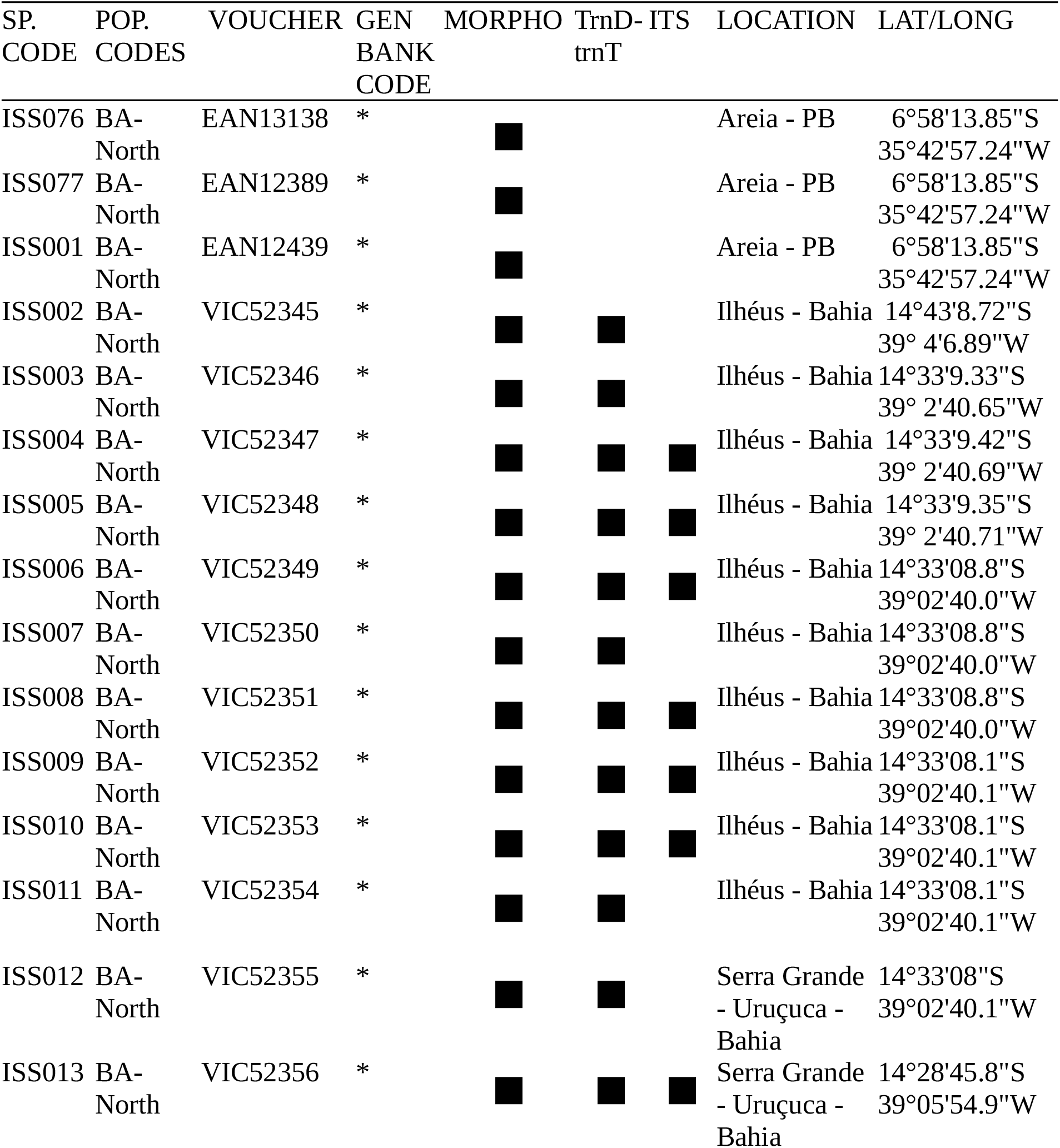
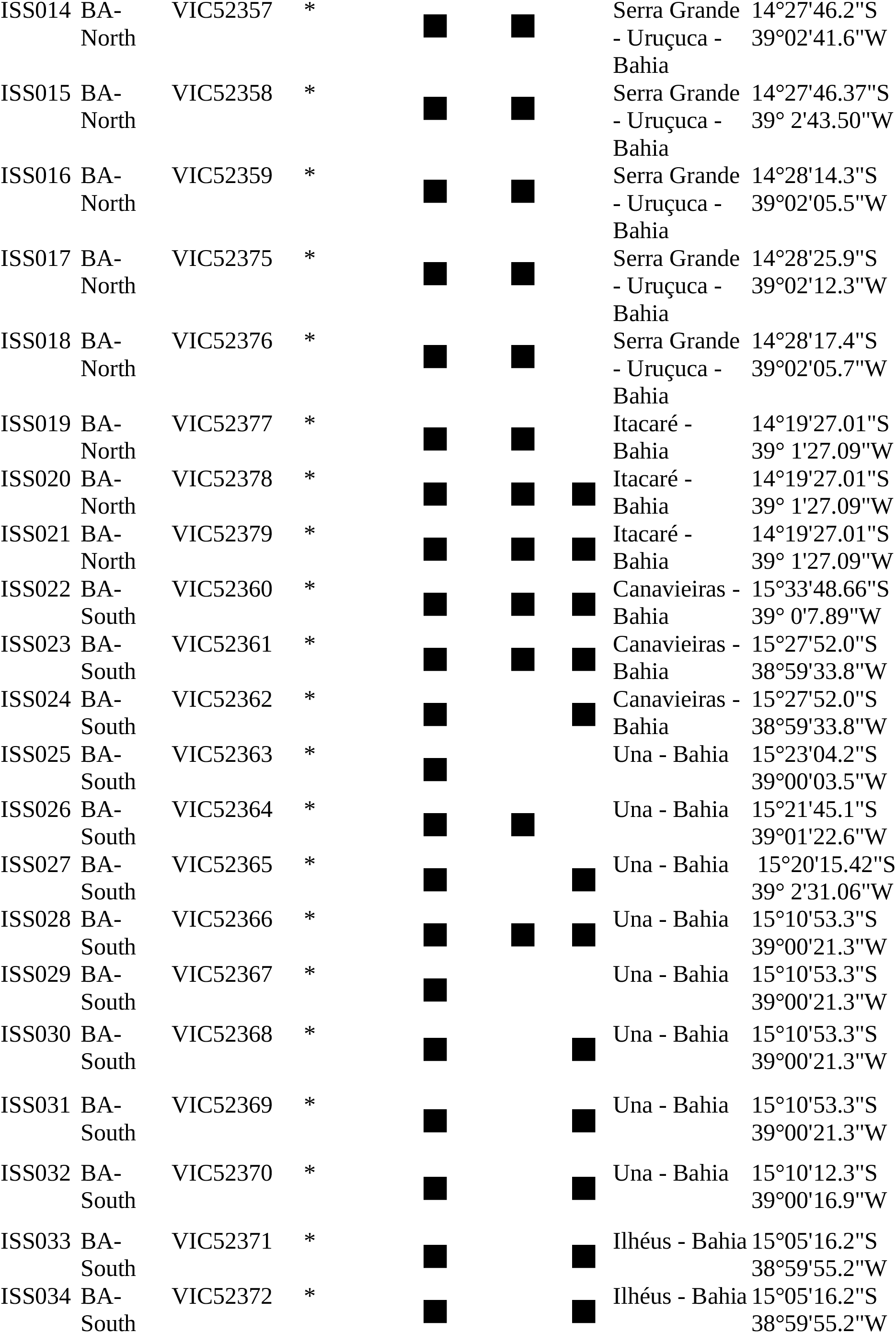
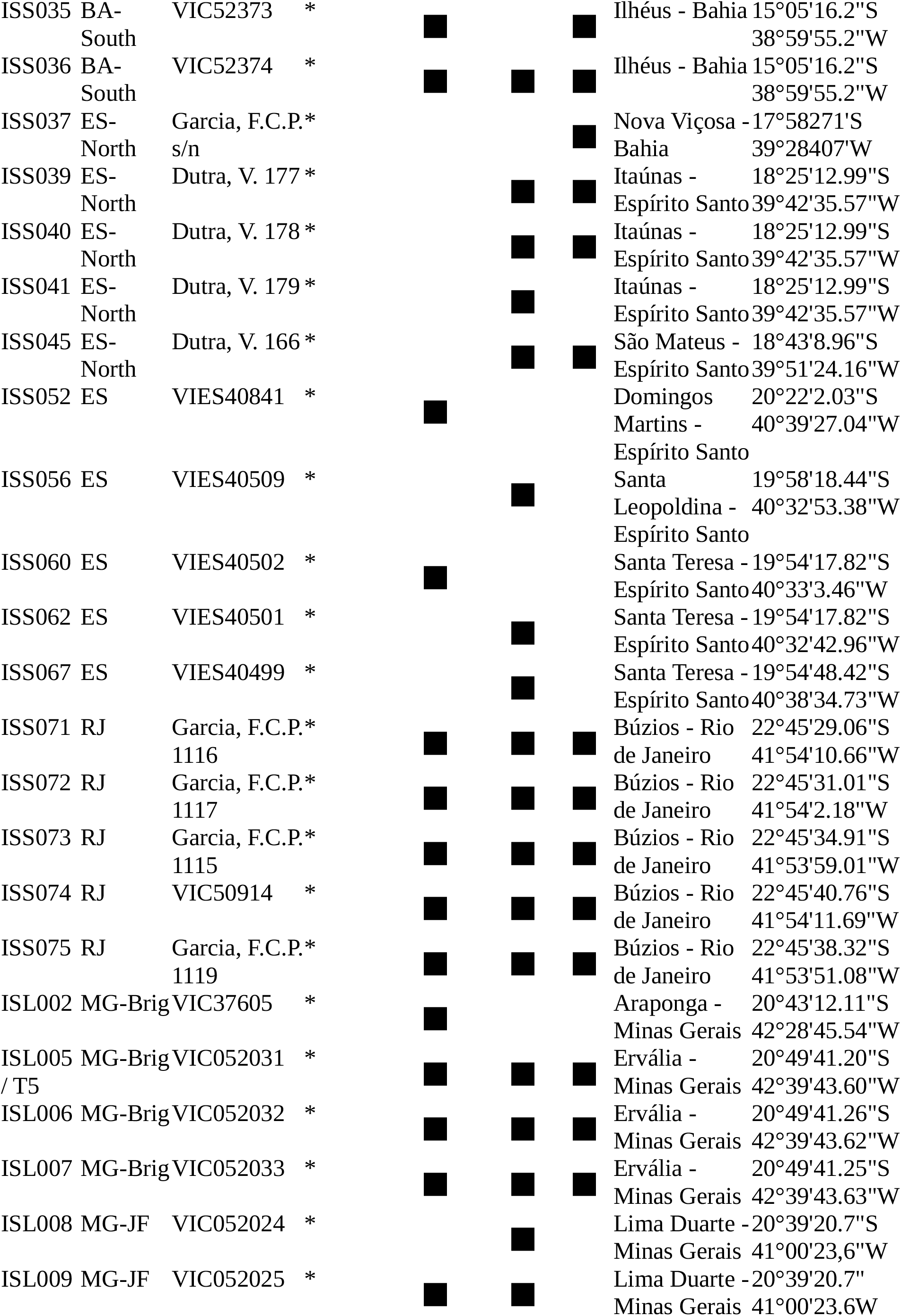
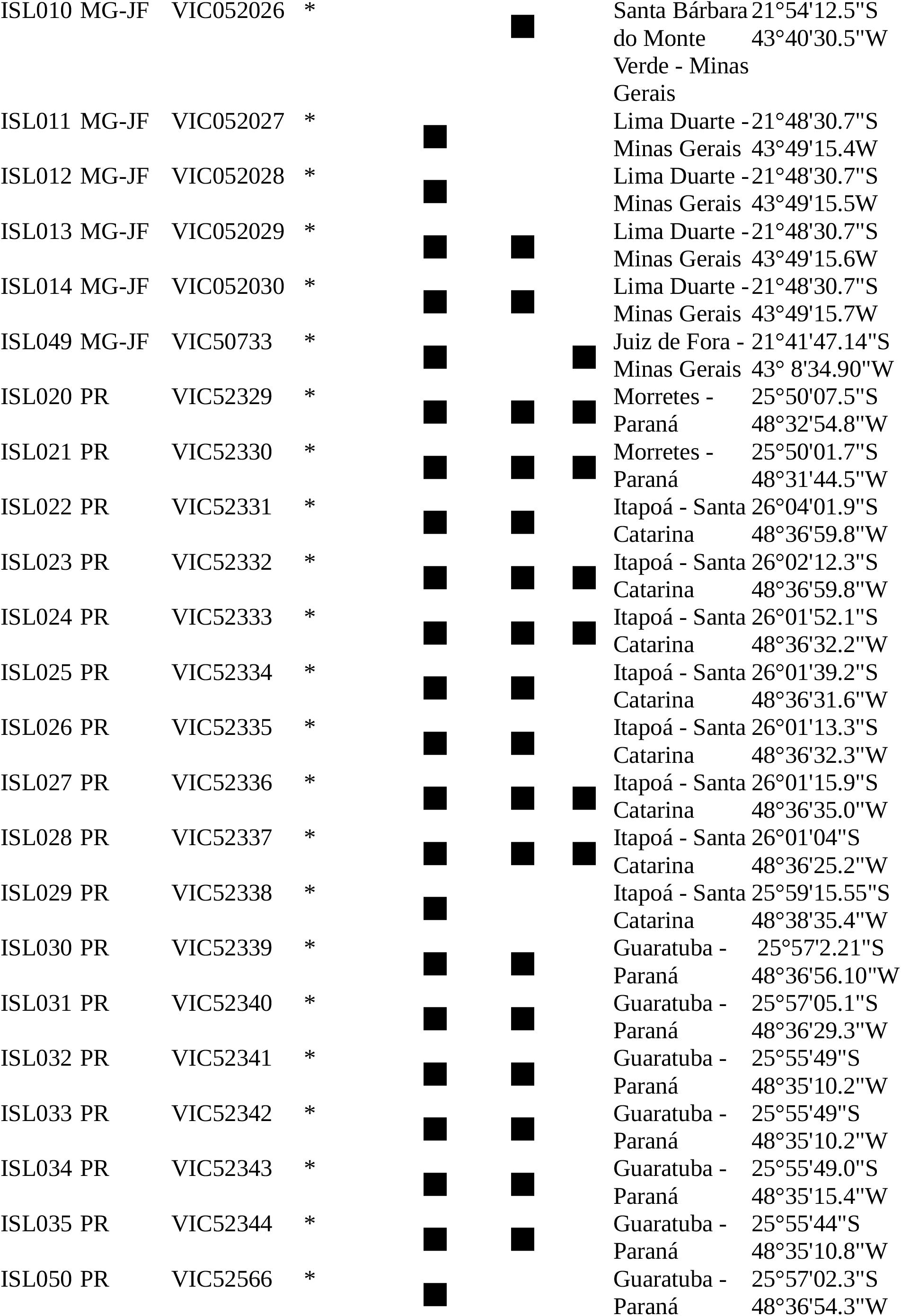
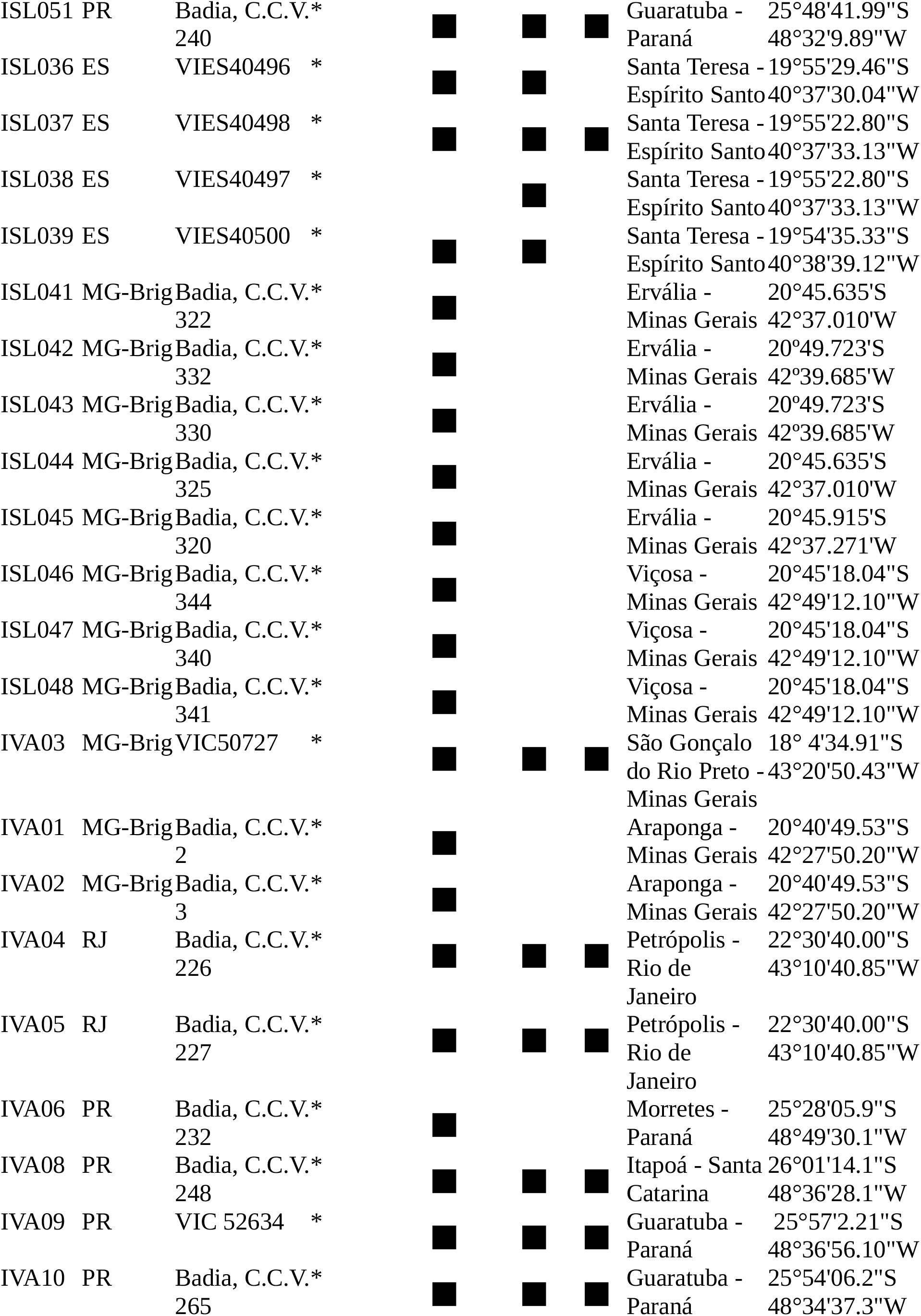
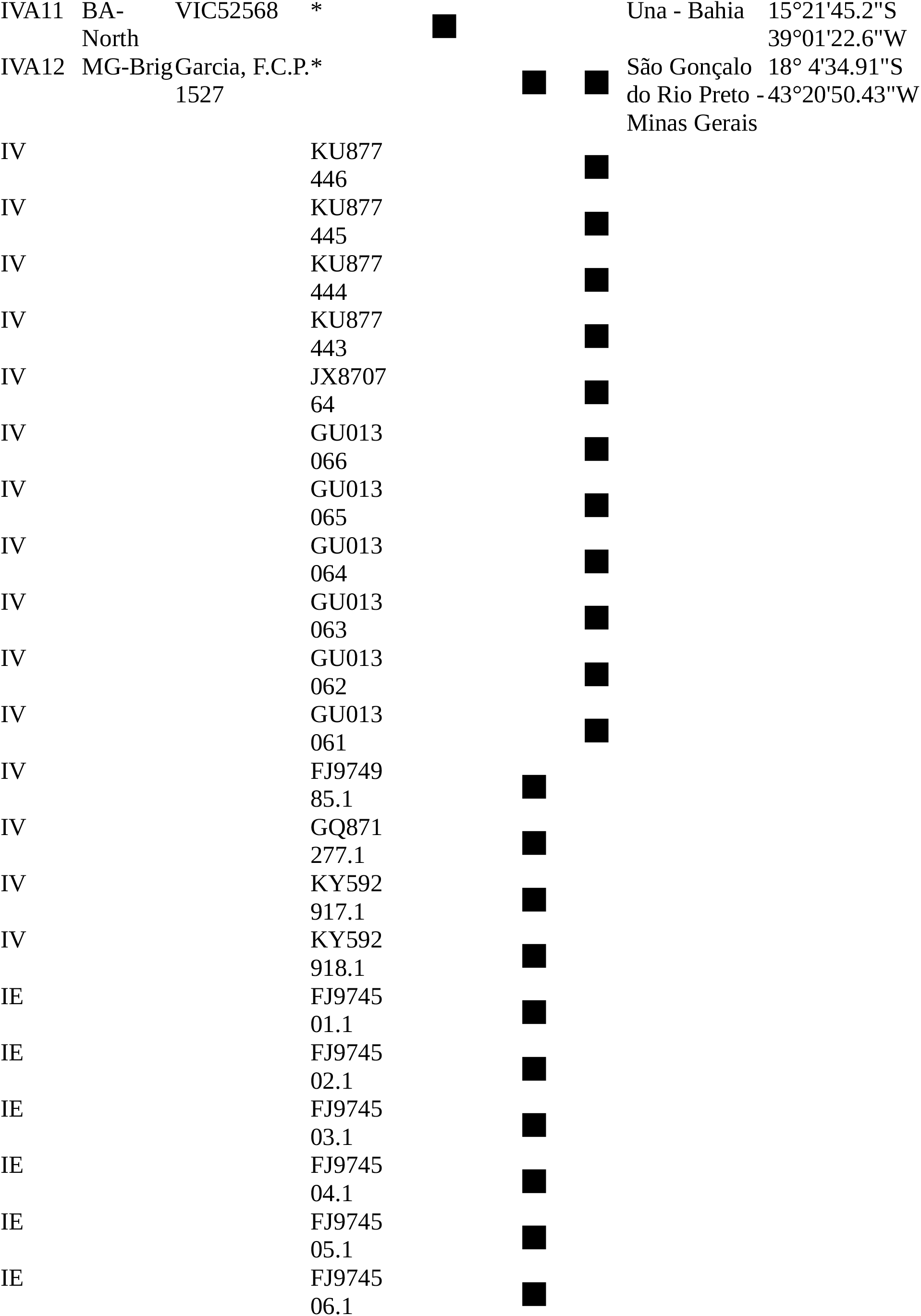
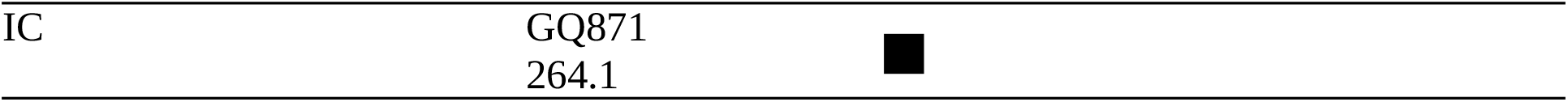
Table providing Specimens Codes (SP. CODE). “ISS” refers to *Inga subnuda* subsp. *subnuda*, “ISL” to *I. subnuda* subsp. *luschnathiana*, “IVA” to *I. vera* subsp. *affinis, “*IV*”* to *Inga vera*, “IE” to *Inga edulis* and “IC” to *Inga capitata*. The population codes (“POP.CODES”), Herbarium Vouchers (“VOUCHER”) Location (“LOCATION”) and Geographical Coordinates (“LAT/LONG” = latitude and longitude) are also provided for the specimens sampled in this work. Sequences retrieved from GenBank (IV, IE and IC codes) were not assigned to any population. Specimens without vouchers are indicated with the collection number. The black squares indicates which methods was applied to which specimen. “MORPHO” = Morphometric Analysis; “D-T” = trnD-trnT analysis; “ITS” = ITS analysis. * indicates samples to be deposited in GenBank.

